# A Unified Protein Embedding Model with Local and Global Structural Sensitivity

**DOI:** 10.1101/2025.10.27.684815

**Authors:** Jerry Xu, Shaojun Pei, Gil Alterovitz

## Abstract

Structural comparison between proteins is key to many research tasks, including evolutionary analysis, peptidomimetics, and functional annotation. Traditional structure alignment tools based on three-dimensional protein structures, such as TM-Align, DALI, or ProBiS, are accurate, but they are computationally expensive and impractical at scale. Existing protein language models (PLMs), such as TM-Vec, improve computational efficiency but only capture global structural similarity, overlooking important motif-level structural details. In this paper, we propose a novel PLM consisting of a Siamese neural network, enabling efficient embedding-based structural comparison while also capturing both global and local structural similarity. Our model was trained on a dual loss function combining TM-score, a global similarity metric, and a variation of lDDT scores, a per-residue similarity metric. We tested against two datasets: a varied TM-score dataset from TM-Vec, and a high TM-score mutant dataset from VIPUR. Against these sets, our model achieved a TM-score MAE of 0.0741 and 0.0583, respectively, and a lDDT-score MAE of 0.0788 and 0.0038, respectively. Our model fulfills two key roles: first, it rapidly detects global structural differences. Second, it supports fine-grained structural assessments, improving sensitivity to subtle but functionally important structural changes.

## 1 Introduction

Many research tasks depend on identifying structural homologs of proteins, including but not limited to evolutionary analysis, peptidomimetics, and functional annotation. However, identifying structural homologs at scale requires efficient comparison algorithms, which currently are still limited.

Over the years, sequence alignment algorithms such as BLAST [1] and MMseqs2 [2] have become extremely optimized, allowing for rapid, large-scale processing. However, sequence alignment alone is ineffective for structural comparisons, since many structural homologs differ vastly in sequence, and likewise many sequential homologs differ vastly in structure [3] [4] [5]. Meanwhile, structural alignment algorithms are computationally expensive due to high algorithmic complexity. In particular, such algorithms often work with *C*_*α*_ distance matrices to directly superimpose proteins, which has a time complexity of *O*(*mn*) or worse for protein pairs of lengths *m* and *n*.

Furthermore, structural alignment algorithms like TM-align [6] and DALI [7] focus on global alignments, even though motif-level or subdomain-level features may be crucial to protein function. Meanwhile, local alignment algorithms like ProBiS [8] only consider surfaces and binding sites, and hence may miss the relevance of internal structural motifs or global folds.

In contrast, protein language models, or PLMs (usually transformer architectures) [9], can predict structural similarity for two proteins of arbitrary length in 𝒪 (1) time given pre-computed embeddings. Utilizing the biophysical prior that sequences alone can reconstruct structures, [10], these models produce fixed-size embeddings directly from sequences that capture structural, chemical, or other features of proteins. Embeddings can then be compared via cosine similarity.

Although efficient, existing PLMs for structural awareness focus on global features rather than local features. One notable example is TM-Vec [11], a recently-developed PLM trained to predict TM-scores (template-modeling scores) [12], a global similarity metric for proteins. While highly accurate for TM-scores, TM-Vec is locally unaware.

Our PLM, consisting of a transformer-based Siamese neural network [13], addresses both the inefficiency of superposition-based algorithms and local insensitivity of prior methods. We continue to generate sequence-based embeddings, resulting in efficient comparison times. Furthermore, our model utilizes a loss function that combines TM-score and a custom variation of lDDT scores (Local Distance Different Test scores) [14], which are per-residue local structural similarity scores, ensuring the model captures both global and local structural features.

The network predicts lDDT scores by producing per-residue embeddings, and predicts TM-scores by pooling those per-reside embeddings into a global embedding. Prior to training, ground truth TM-scores and sequence alignments are computed using TM-align, and ground truth lDDT scores are computed by comparing the local atomic environments of *C*_*α*_ atoms in aligned residues. During training, predicted TM-scores are calculated as the cosine similarity between the global embeddings of the two proteins. Using the sequence alignment generated by TM-align, we predict the lDDT scores for each alignment pair as the cosine similarity of the embeddings for the residues in that pair.

Our paper’s main contribution is the generation of protein language embeddings which are both locally and globally structure-aware. These embeddings can then be used for efficient structural comparison in many downstream research tasks, particularly in mutation analysis. Global embeddings can be utilized to determine the degree to which the overall fold of the mutant differs from the wild-type protein, quickly identifying deleterious mutants (i.e. low TM-score mutants). Per-residue embeddings can further analyze these mutants, identifying affected subdomains and hence the impacted functions of the protein.

## 2 Methods and Materials

Our Siamese network-based architecture consists of three main steps.

- **Data Preprocessing**: The purpose of this step is twofold. Firstly, it generates the ProtTrans embeddings for each protein that will be the inputs of the neural network. Secondly, it precomputes the true TM-scores and lDDT scores, as well as the sequence alignments used to predict lDDT scores. All ground truth information is cached.
- **Siamese Neural Network**: For every pair of proteins whose similarity will be predicted, the (padded) ProtTrans embeddings of both proteins will be independently processed by a neural network with shared weights. The transformer module will capture sequential dependencies, as well as local and global structural patterns, producing learned per-residue embeddings that contain local structural information for use in lDDT score prediction. After being processed by the transformer module, the per-residue embeddings will be pooled via an attention mechanism to produce a single global embedding for each protein, to be used TM-score prediction.
- **Contrastive Loss**: The per-residue lDDT score loss and TM-score loss are combined for backpropagation. The degree to which each type of loss contributes to model learning depends on the true TM-score of that training pair.

### 2.1 Data Preprocessing

#### 2.1.1 Datasets

This study utilized datasets from UniProt and SWISS-MODEL, particularly working with protein sequences and PDBs.

For training, our model used a dataset from SWISS-MODEL consisting of 250, 000 distinct proteins with length at most 300 amino acids, organized into 300, 000 protein pairs. This dataset mimics the training dataset of TM-Vec [11], which can be can be found in full at https://zenodo.org/records/8038377 in the file swiss under 300 141M.csv. PDBs were obtained from SWISS-MODEL’s REST API at https://swissmodel.expasy.org/repository/uniprot/<id>.pdb?provider=swissmodel.

For testing, two benchmark datasets were used:

- **TM-Vec dataset**: For this dataset, we reserved 886 protein pairs from the SWISS-MODEL dataset that were not used in training.
- **VIPUR dataset**: This dataset consists of 350 curated human wild-type/mutant pairs, some benign and some deleterious, from the dataset used in VIPUR, a model trained to predict deleteriousness of protein variants [15].

These datasets were designed to test the robustness of the model against both global structural similarity and local structural similarity.

For each protein in the TM-Vec dataset and the wild-type proteins in the VIPUR dataset, PDBs were downloaded based on the AlphaFold-generated model of the proteins using the AlphaFold PDB API at https://alphafold.ebi.ac.uk/files/AF-<uniprot-id>-F1-model_v4.pdb. (This might represent a slight discrepancy from how TM-Vec evaluated their results, as they may not have used AlphaFold PDBs.)

In order to induce the desired structural changes for each mutation in the VIPUR dataset, we used the `MODELLER` library with a sphere size of 100Å to remodel the AlphaFold PDB of the wild-type protein. Figure 1 provides a visual overview of the changes induced by `MODELLER` for one protein, the human tumor suppressor ARF.

**Figure 1:**
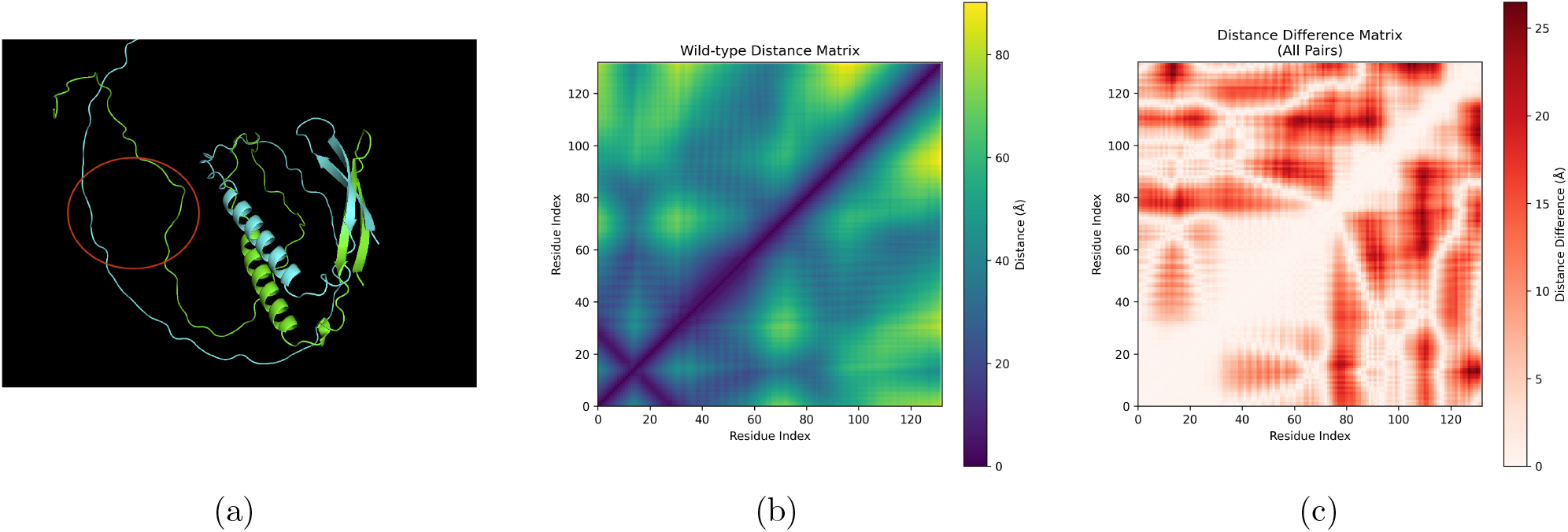
MODELLER’s remodeling for the human tumor suppressor ARF (UniProt ID Q8N726). (a) The wild-type and mutant PDB superimposed, with the mutation site highlighted in red. (b) The distance matrix between residues in the wild-type protein. Certain structural features can be identified: the purple line parallel to the main antidiagonal around residue 30 represents the beta hairpin at the start of the protein, and the thickened section of purple around residues 40-60 along the main diagonal represents the alpha helix after the hairpin. (c) The difference matrix of the distance between two residues in the wild-type PDB and remodeled PDB. Darker red colors close to the main diagonal are indicative of local structural changes, while darker red colors farther from the main diagonal are indicative of global structural changes.

#### 2.1.2 Sequence Embeddings

We utilize the pre-trained ProtT5-XL-UniRef50 model [16] to generate per-residue embeddings of length 1024 for each amino acid in the protein sequence. Since the proteins in the datasets have variable lengths, each protein’s per-residue embeddings are padded before training such that there are 300 per-residue embeddings (the maximum sequence length), simultaneously producing a binary padding mask that indicates which embeddings are padded. Hence, the per-residue embeddings for the whole protein satisfy **X** ∈ ℝ^300*×*1024^.

#### 2.1.3 TM-scores

TM-score, or **t**emplate **m**odeling score, is a commonly-used metric of global structural similarity in proteins that is normalized between 0 and 1, with higher scores indicating higher global similarity [12]. TM-score depends on a sequence alignment between the reference and model proteins, and is calculated as

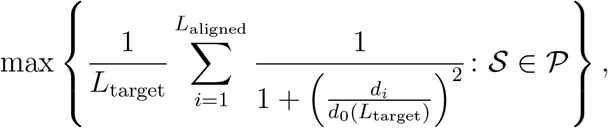

where 𝒫 is the set of superpositions of the template and target structures, *L*_target_ is the number of residues of the target structure, *L*_aligned_ is the number of residues aligned between the template and target sequences, *d*_*𝒮*_ (*i*) is the distance between the *i*th pair of aligned residues in the template and target sequences in 𝒮, and 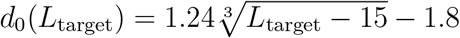 Here, each residue is represented as the point corresponding to the position of its *C*_*α*_.

#### 2.1.4 TM-align

TM-align is a superposition-based algorithm for global structural alignment [6]. We use the `tm_align` function in the `tmtools` python library to extract two pieces of ground truth information: first, we obtain true TM-score (`results.tm_norm_chain1`) between the two proteins (we normalize the TM-score against chain1, the reference protein). Second, we obtain the residue alignment information (`seqxA, seqM, seqyA`) we can use to calculate predicted lDDT scores, which is given in the below format:

~~~
seqxA: GKTIQVIPHVTNEIKDFISIGED—EVDFMLCEIG
seqM: ::::::::::: :::.::: ::: ::::::::::
seqyA: GATVQVIPHVT-ALKEKIKRAATTTDSDVIITEVG
~~~

“:” denotes highly aligned residues, “.” denotes poorly aligned residues, and “” (space) denotes unaligned residues. “-” represents gaps in one of the sequences.

#### 2.1.5 lDDT Scores

lDDT scores, which stands for **l**ocal **D**istance **D**ifference **T**est scores, were initially used to determine how well a predicted (target) structure of a protein matched the experimentally-determined (template) structure of that protein [14]. Per-residue lDDT scores are also normalized between 0 and 1 and assigned to each amino acid in the target protein, representing how well the local atomic environment of that residue matches the local atomic environment of the same residue in the template protein (with higher scores meaning more aligned). The original algorithm for lDDT scores is described in Algorithm 1.

##### Algorithm 1 Per-Residue lDDT Scores

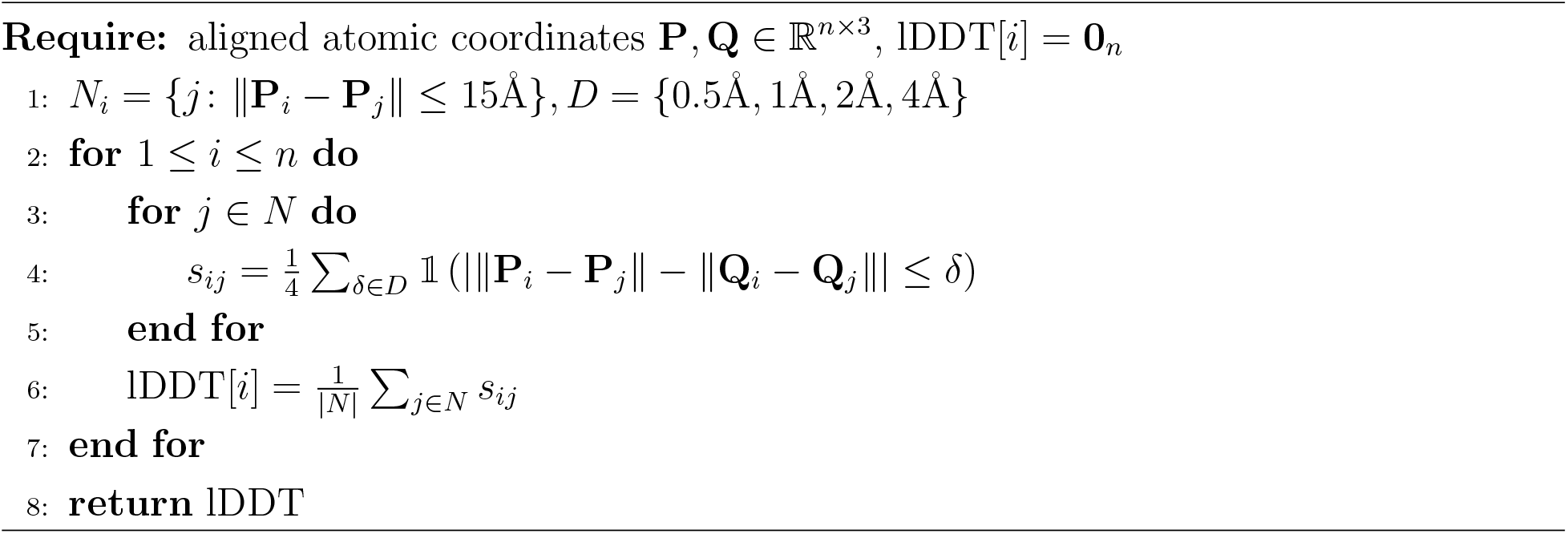

We cannot directly use this implementation of lDDT scores, since it depends on the two proteins having identical sequences and hence an atomic bijection. In our case, there is no longer a clear one-to-one sequential alignment between proteins, and aligned residues are also unlikely to have an equal number of atoms. Therefore, we made the following adjustments:

- First, we align residues using TM-align’s alignment information, and generate a list of aligned residue pairs: all unaligned residues are discarded from the rest of the calculation. To resolve the lack of an atomic bijection, we only use the *C*_*α*_ atoms of each residue in an alignment pair. This means that only inter-*C*_*α*_ distances are compared.
- We calculated the per-residue lDDT scores using protein 1 as the template and protein 2 as the target. Furthermore, the array of lDDT scores was padded to length 300: each aligned residue will have its respective lDDT score inputted in the appropriate position in the array, while all non-aligned or padded residues were assigned an lDDT score of 0.
- We maintained the neighborhood threshold of 15Å and the difference thresholds of *{*0.5Å, 1Å, 2Å, 4Å*}*. However, since there are far fewer *C*_*α*_ atoms within this neighborhood than total atoms, we added a distance-decaying kernel to ensure that closer distances are upweighted in their contribution to lDDT score. After experimenting with different kernel types, we found that an inverse cubic kernel provided the most stable lDDT scores while still highlighting important local structural changes.

The revised algorithm is given in Algorithm 2.

##### Algorithm 2 Custom Per-Residue lDDT Scores

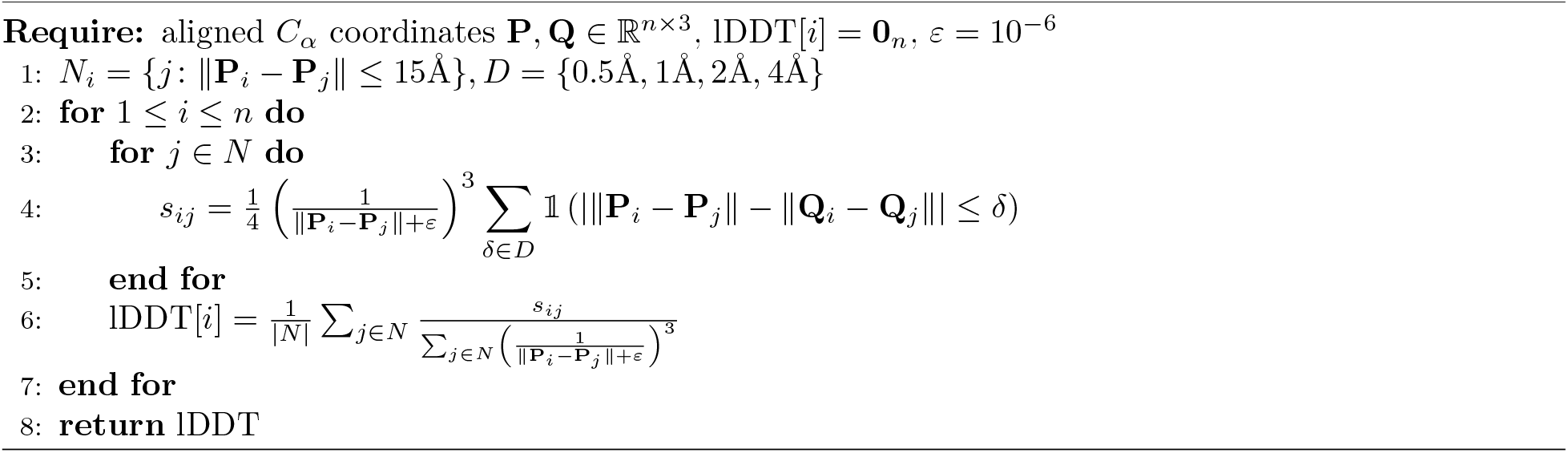

### 2.2 Siamese Neural Network

Siamese neural networks are used for tasks involving pairwise comparison and similarity analysis, and hence take in two inputs. The typical architecture of a Siamese neural network consists of a feature extractor and a comparison head, as illustrated in Figure 2. The feature extractor consists of two branches *f*_1_ and *f*_2_, which have the same network structure and weights (*W*) [13]. The comparison head outputs a comparison score *C*_score_ based on the features *f*_1_(*x*_1_) and *f*_2_(*x*_2_) extracted from the inputs *x*_1_ and *x*_2_, respectively.

**Figure 2:**
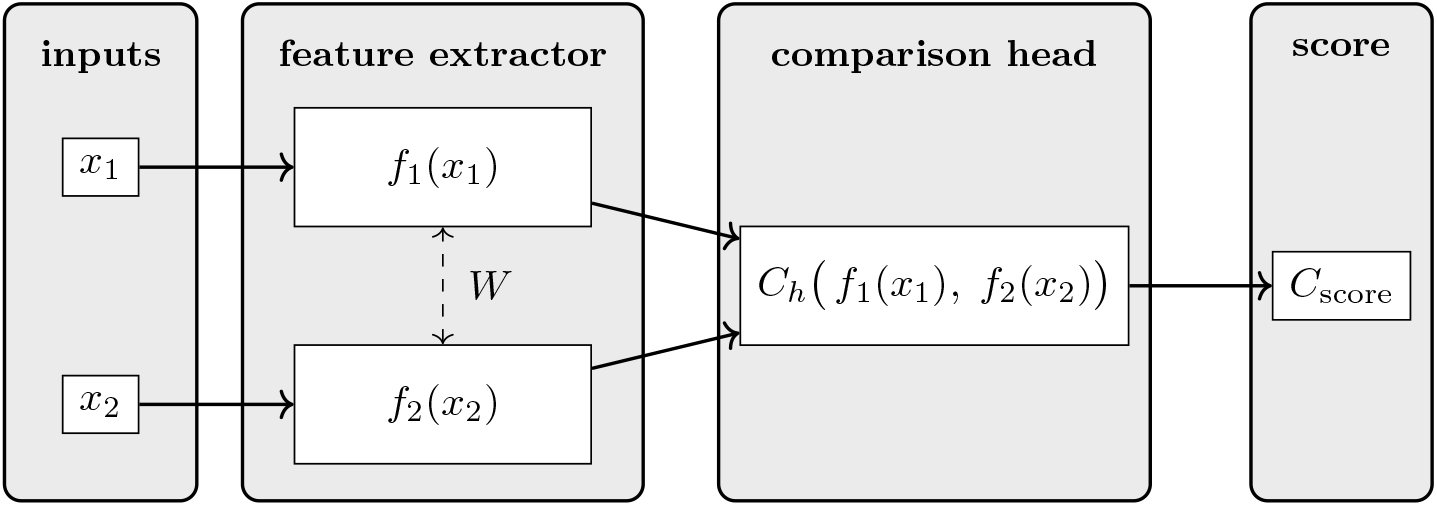
The typical architecture of a Siamese neural network consists of a feature extractor (i.e. the two neural networks, with shared weights) and a comparison head.

There are generally two comparison mechanisms: similarity comparison, which calculates a form of distance between the feature vectors, and ranking comparison, which orders elements based on some metric.

Our feature extractor consists of the pretrained ProTrans model, an input projection, two transformer layers, pooling, and an output projection (shown below). Our Siamese neural network utilizes similarity comparison.

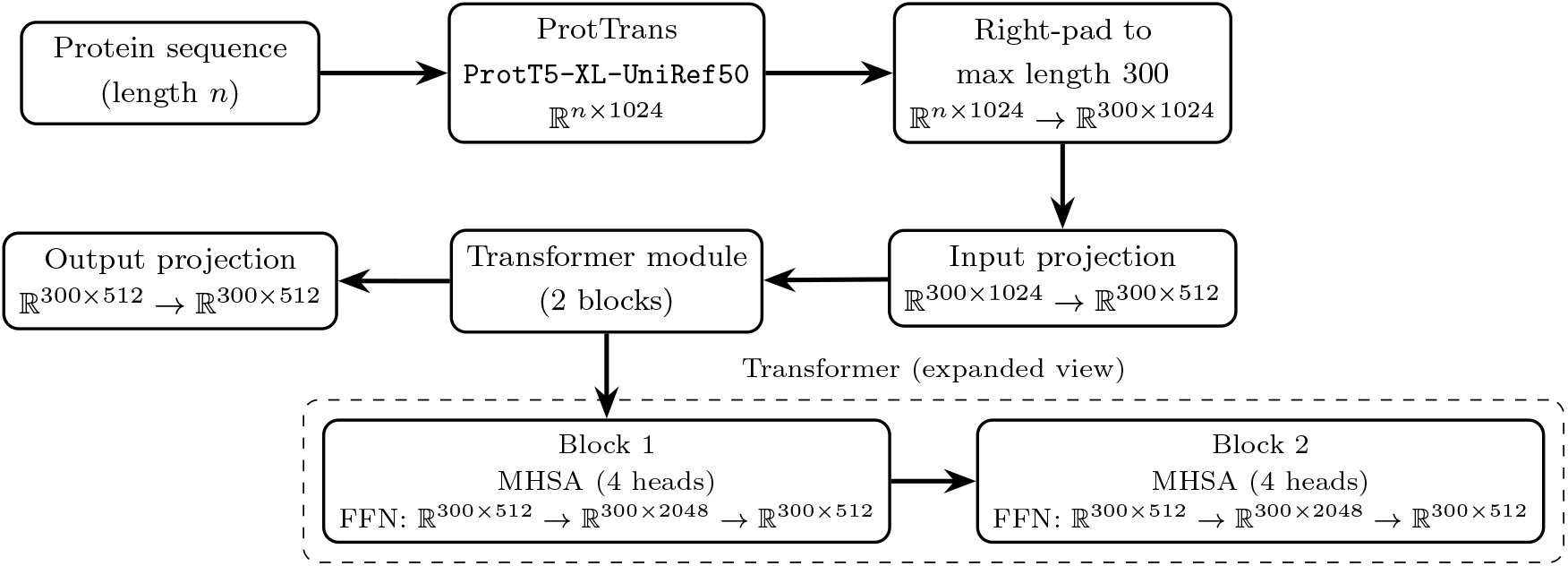

#### 2.2.1 Inputs

The two neural networks take in the padded embeddings from ProtTrans of the form **X** ∈ ℝ^300*×*1024^, as well as the padding mask of shape **M** ∈ ℝ^300^. Before passing through the transformer, the model must pass through an input projection consisting of a linear layer, layer normalization [17], and Bernoulli dropout [18].

#### 2.2.2 Transformer Module

Each neural network is composed of two transformer encoder layers and an attention pooling mechanism. The transformer layers consist of four attention heads and two feedforward layers each. Each multi-head attention component runs scaled dot-product attention several times in parallel, as described in Algorithm 3. The feedforward layers consist of two linear layers using ReLU as an activation function, layer normalization, and Bernoulli dropout.

##### Algorithm 3 Multi-Head Attention

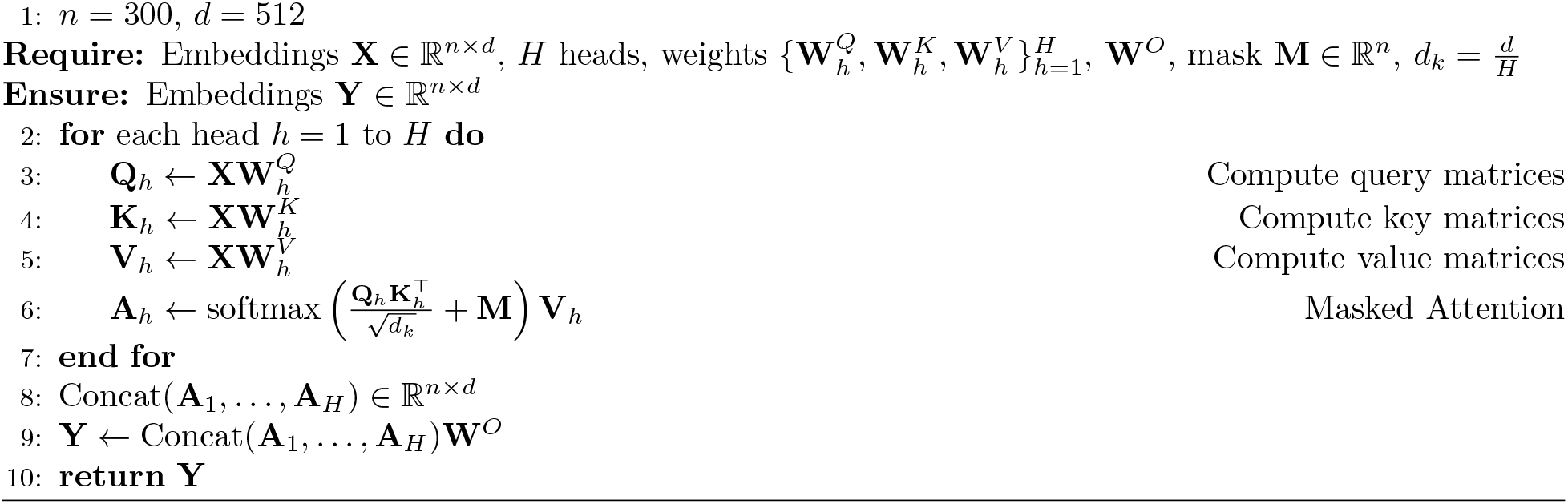

#### 2.2.3 Pooling

We implement softmax pooling to produce a single, structurally-aware embedding representing the protein, which is used in TM-score prediction. An attention layer first assigns weights to the per-residue embeddings using softmax normalization. Then, the global embedding is created by computing a weighted average of the per-residue embeddings. This is described in Algorithm 4.

##### Algorithm 4 Attention Pooling

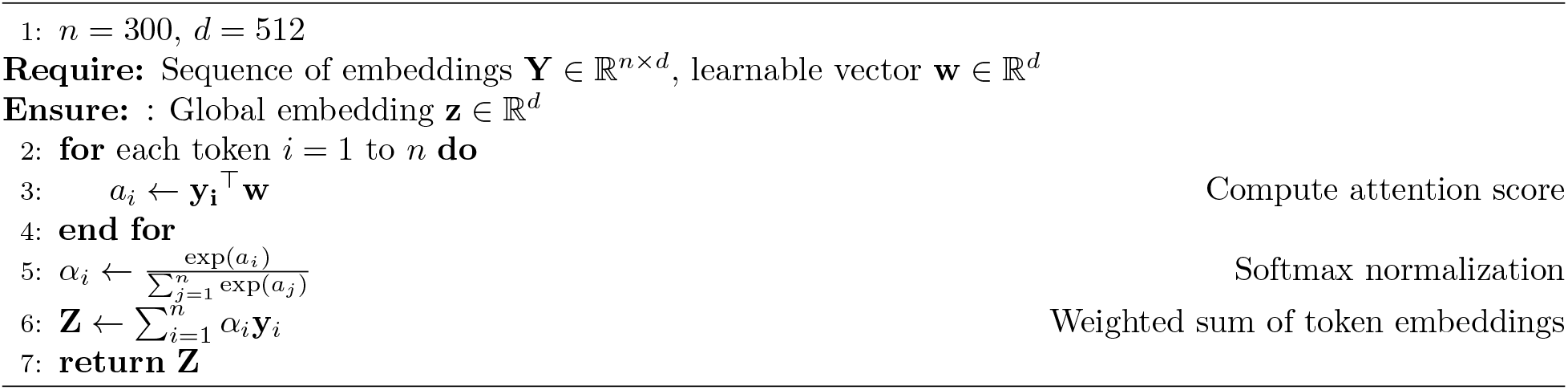

#### 2.2.4 Output

The outputs of each neural network are passed through an output projection consisting of a linear layer using ReLU as an activation function. The final per-residue embeddings are of the form **Y** ∈ ℝ^300*×*512^, and the final global embedding is of the form **Z** ∈ ℝ^512^.

### 2.3 Contrastive Loss

Contrastive loss was based upon a combined loss function involving both TM-score (allowing for global structural sensitivity) and lDDT score (allowing for local structural sensitivity). Pairs with true TM-scores less than 0.1 were ignored since their structures are too dissimilar to extract useful patterns. Pairs with true TM-scores less than 0.7 utilized only TM-score loss since local structural similarity requires sufficient global alignment between proteins. Pairs with true TM-scores at least 0.7 used both lDDT and TM-score loss. The specific loss function we used is detailed below:

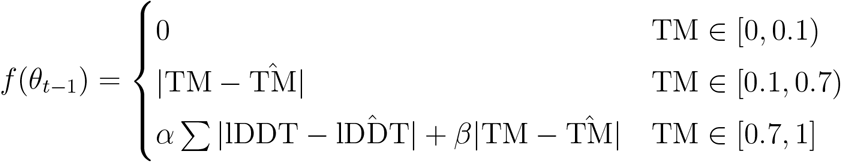

(In our model, *α* = 0.7, *β* = 0.3. *θ*_*t*_ are the parameters at time *t*.)

Predicted TM-scores and lDDT scores were computed using cosine similarity, and weights were updated using Adam optimizers [19].

- Predicted per-residue lDDT values were calculated from the per-residue embeddings using TM-align’s sequence alignments. Given residue *i* in protein 1 is aligned with residue *j* in protein 2, with respective embeddings 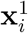 and 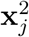, then 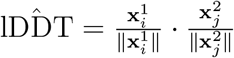.
- Predicted TM-scores were calculated directly from the pooled global embeddings **z**_1_ and **z**_2_ for proteins 1 and 2 as 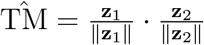.

Figure 3 shows the architecture of one of the two neural networks, including the subsequent prediction and loss steps. (Projections are omitted from this diagram.)

**Figure 3:**
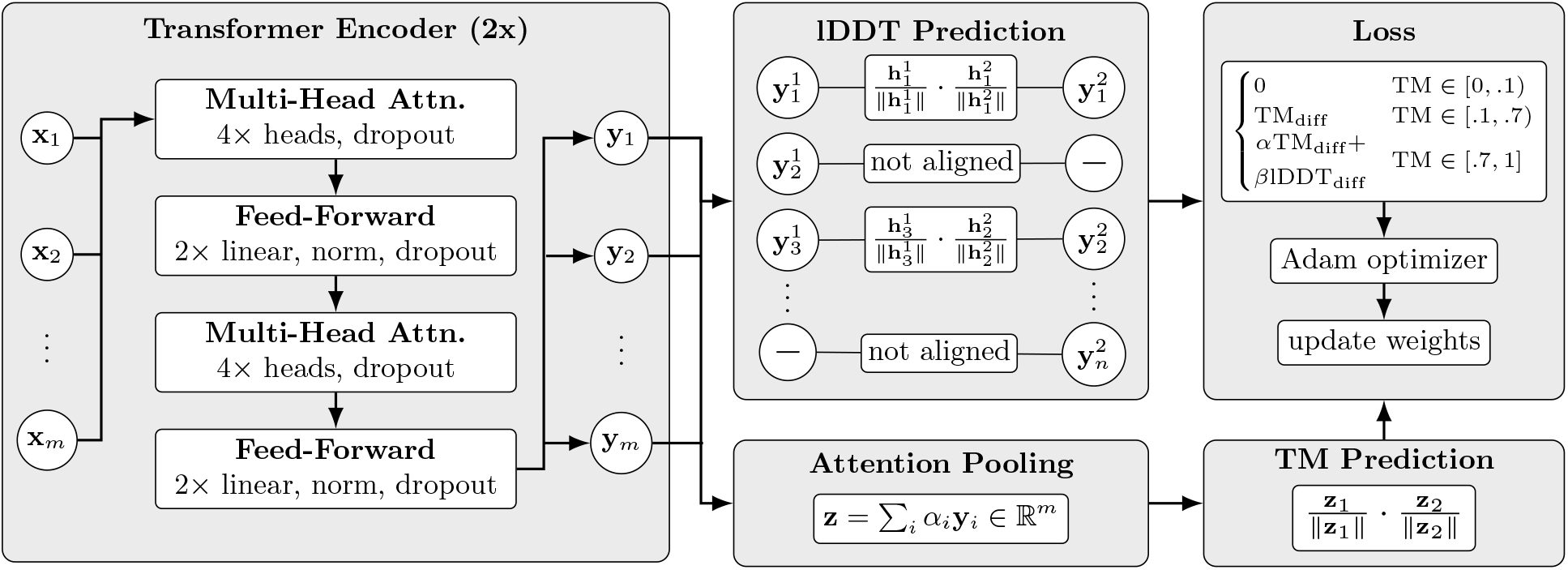
The structure of a single neural network. There are two transformer encoder layers with four attention heads and two feedforward layers each. The per-residue embeddings outputted by the transformer layers are used to generate predicted lDDT scores via cosine similarity using the sequence alignment generated by TM-align. After being processed by the transformer, the per-residue embeddings are pooled into a global embedding that generates a predicted TM-score via cosine similarity. The loss function uses both TM-score and lDDT scores.

### 2.4 Training

Training occurred over 5 epochs of the SWISSS-MODEL dataset. The parquet files were shuffled prior to each epoch to prevent overfitting. The batch size was 16 pairs. V100 GPUs from Pittsburgh Supercomputing Center’s Bridges-2 were used to accelerate the speed of training.

### 2.5 Comparison to Prior Work

Here, we compare our framework with other well-known structural prediction tools. The differences are summarized in Table 1. All time complexities are given for the task of comparing two proteins of lengths *m* and *n*.

**Table 1:** Comparison of our model’s framework against selected PLMs.

| Tool | Framework | Time Complexity | Predicts |
| --- | --- | --- | --- |
| **TM-Vec** | Transformer-based Siamese neural network | $\mathcal{O}(1)$ | Global structural similarity |
| **DALI** | Superposition-based global alignment tool | $\mathcal{O}(m^2n^2)$ | Global structural alignment |
| **TM-align** | Superposition-based global alignment tool | $\mathcal{O}(mn)$ | Global structural alignment and similarity |
| **ProBiS** | Graph-based local surface alignment tool | exponential | Locally similar surfaces/binding sites |
| **Our model** | Transformer-based Siamese neural network | $\mathcal{O}(1)$ | Global and local structural similarity |

Hamamsy et al. (2023) [11] designed TM-Vec, which is also a PLM in the form of a transformer-based Siamese neural network. Hence, it has a time complexity of *O*(1). In Hamamsy’s paper, TM-Vec was used in conjunction with DeepBLAST, a structural alignment algorithm. TM-Vec predicts the structural homologs of a query protein within a vector database, which are then passed into DeepBLAST. DeepBLAST utilizes differential sequence alignment algorithms to predict areas of local structural alignment in those homologs. Unlike our model, TM-Vec is only trained to predict TM-scores. We will be using TM-Vec as a baseline to evaluate the performance of our model’s TM-score prediction.

Holm et al. (1993) [7] created DALI, a superposition-based global alignment algorithm. DALI creates a *C*_*α*_ distance matrix for both proteins, and attempts to string together local alignments of 6 *×* 6 submatrices (corresponding to hexapeptide fragments) to form a global alignment. Each pair of fragments is assigned a similarity score that indicates how well the submatrices align. A graph is constructed using residues as nodes and strong fragment pairs as edges, slowly building up a global alignment. The algorithm utilizes Monte Carlo optimization [20], and is hence stochastic. Unlike our PLM, which utilizes sequences as input, DALI utilizes coordinates as input. Furthermore, because it must compare the distance matrices of both proteins, its time complexity is 𝒪 (*m*^2^*n*^2^), much slower than our model’s 𝒪 (1).

Zhang et al. (2005) [6] developed TM-align, another superposition-based global alignment algorithm. This algorithm works similarly to DALI, first generating multiple possible initial alignments by matching local fragments (essentially contact patterns, small-scale *C*_*α*_ distance matrices) and secondary structures. Unlike DALI, TM-align uses dynamic programming, allowing the algorithm to try attaching different local fragments onto the current alignment. For each possible alignment, the optimal superposition is computed using the Kabsch-Umeyama algorithm [21]. Scoring matrices (essentially *C*_*α*_ distance matrices in each alignment) are computed, and the alignment maximizing the TM-score is chosen as the final alignment. Like DALI, TM-align also utilizes coordinates as an input. However, this method was much faster than DALI for two reasons: first, it was deterministic, and second, the Kabsch-Umeyama algorithm utilizes singular value decomposition, which as a time complexity of 𝒪 (1). Despite this, the score matrix generation and dynamic programming steps still have a time complexity of 𝒪 (*mn*), resulting in an overall time complexity of 𝒪 (*mn*). This is slower than our PLM’s 𝒪 (1).

Konc et al. (2010) [8] developed ProBiS, a tool for identifying structurally similar protein surfaces or binding sites. ProBiS represents the solvent-accessible surface of each protein as a 3D graph, and uses a maximum clique algorithm to find maximal common subgraphs (MCS) between two proteins. The MCS is used to generate superpositions. A maximum clique search theoretically has an exponential time complexity (in fact, the most efficient maximum clique algorithm has a time complexity of 𝒪 (2^*n/*4^) [22]), although the practical runtime for this algorithm is typically near-polynomial. At the very least, creation of the 3D graph is an 𝒪 (*n*^2^) process. This is much slower than our PLM. Furthermore, ProBiS does not capture similarity in the global fold or internal structural features, which can often be important, while our model addresses both global and local similarity at every region of the protein.

## 3 Results

We evaluated the performance of our model to predict both TM-score for protein pairs and lDDT score for aligned residues against the two testing datasets mentioned in 2.1.1, comparing our TM-score prediction capability against TM-Vec.

### 3.1 TM-Vec Dataset

For the 886 proteins from the TM-Vec dataset, our performance is evaluated against TM-Vec’s in Table 2. The error plots are shown in Figure 4. In Figure 5, we plot both our model and TM-Vec’s performance at each TM-score interval.

**Table 2:** Performance of our model and TM-Vec on the TM-Vec dataset.

| Metric | Model | MAE | MSE | Error Stdev |
| --- | --- | --- | --- | --- |
| IDDT (per-residue) | Our model | 0.0788 | 0.0344 | 0.1224 |
| TM (per-pair) | Our model | 0.0741 | 0.0103 | 0.1010 |
| TM (per-pair) | TM-Vec | 0.0792 | 0.0126 | 0.1113 |

**Figure 4:**
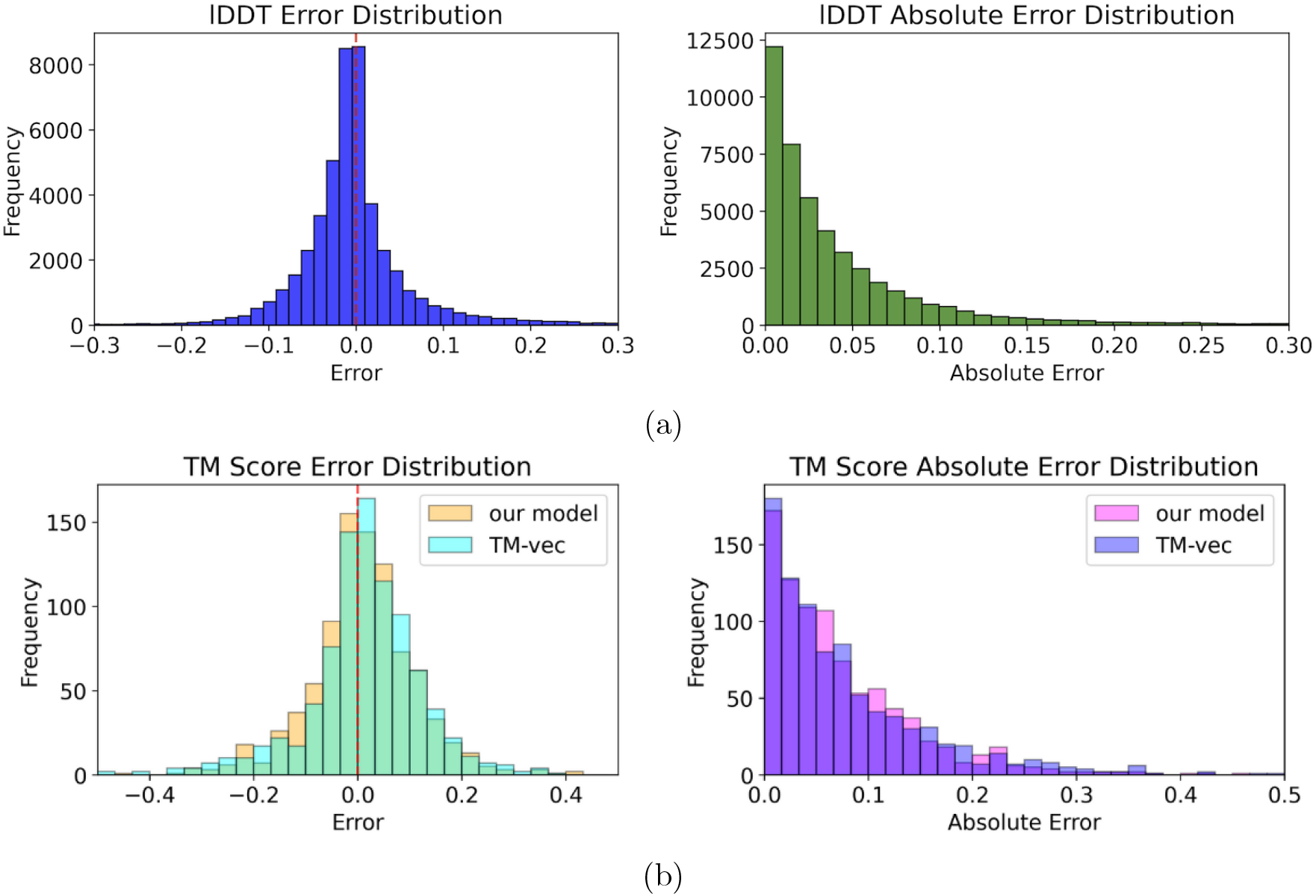
(a) The lDDT score error histograms across all residues. (b) The TM-score error histograms across all protein pairs for both our model and TM-Vec.

**Figure 5:**
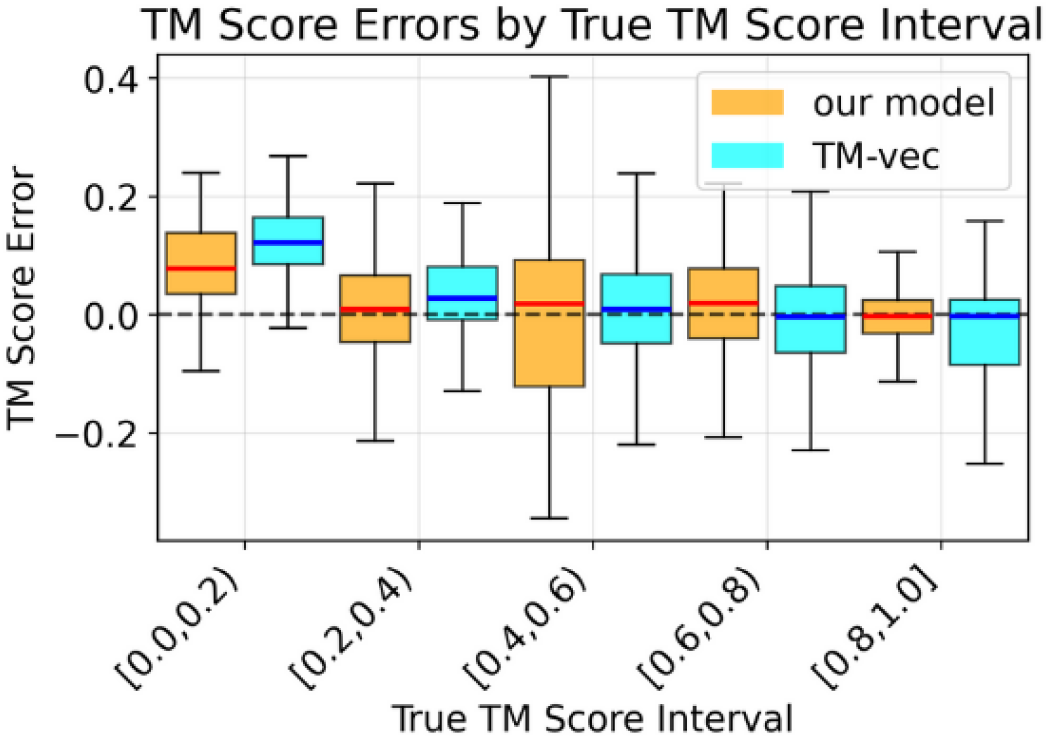
The box plots for our model and TM-Vec’s errors at different TM-score intervals.

When performing Welch’s t-test for our model’s TM-score prediction and TM-Vec’s TM-score prediction with *α* = 0.05, we obtain the following test statistic and degrees of freedom:

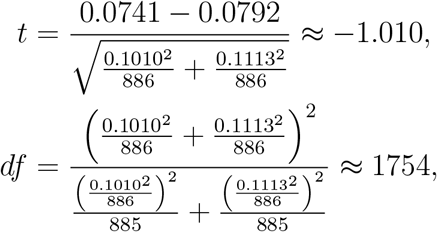

giving *p* ≈ 0.313 *> α*. Thus, across the TM-Vec dataset, there is no statistically significant difference between our model’s and TM-Vec’s TM-score prediction.

### 3.2 VIPUR Dataset

For the 350 proteins in the VIPUR dataset, our accuracy is evaluated against TM-Vec’s in Table 3. The error plots are shown in Figure 6.

**Table 3:** Performance of our model and TM-Vec on the VIPUR dataset.

| Metric | Model | MAE | MSE | Error Stdev |
| --- | --- | --- | --- | --- |
| IDDT (per-residue) | Our model | 0.0038 | 0.0001 | 0.0095 |
| TM (per-mutant) | Our model | 0.0583 | 0.0096 | 0.0980 |
| TM (per-mutant) | TM-Vec | 0.0617 | 0.0102 | 0.1011 |

**Figure 6:**
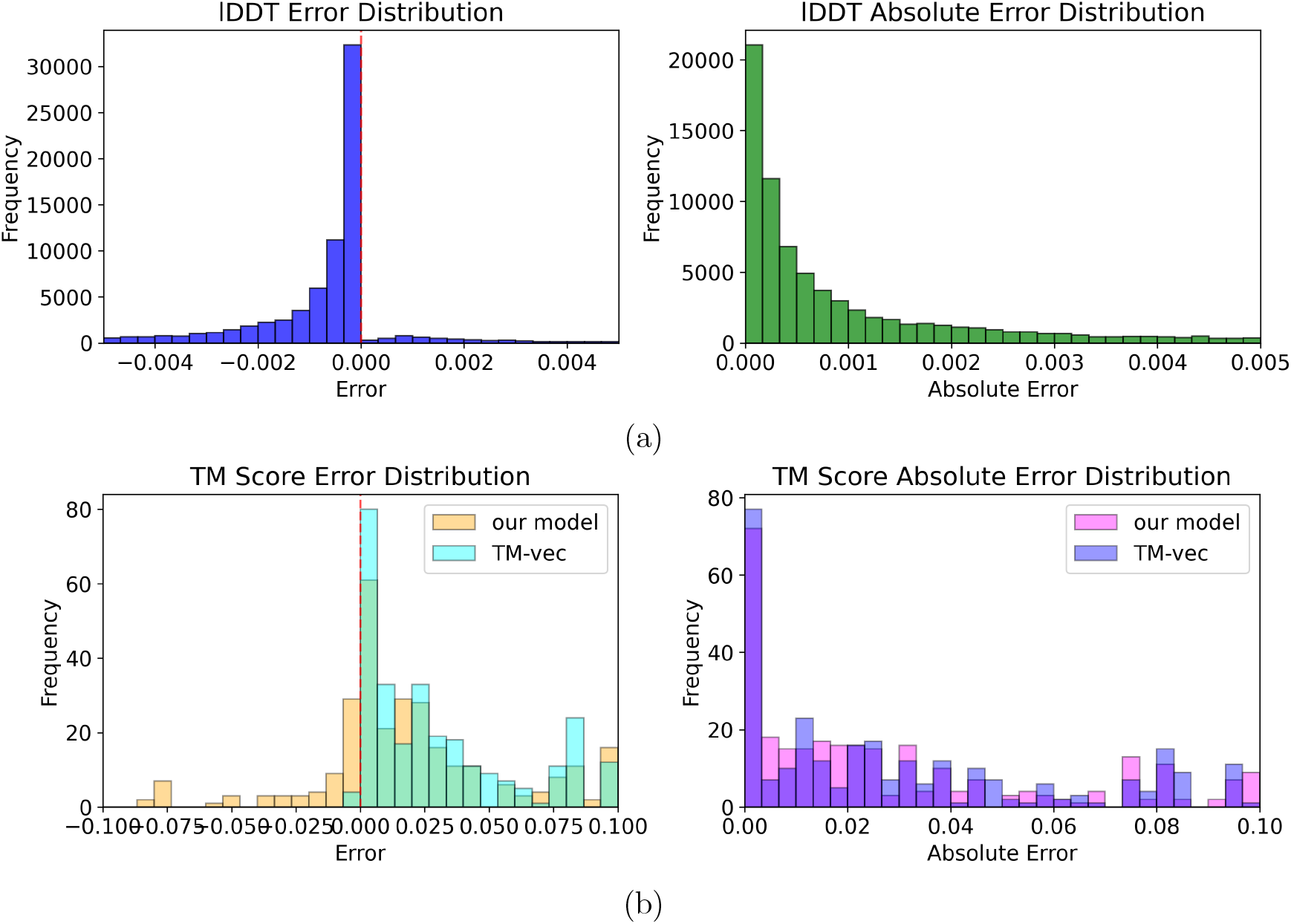
(a) The lDDT score error histograms across all residues. (b) The TM-score error histograms across all mutants for both our model and TM-Vec.

Once again performing Welch’s t-test, we obtain

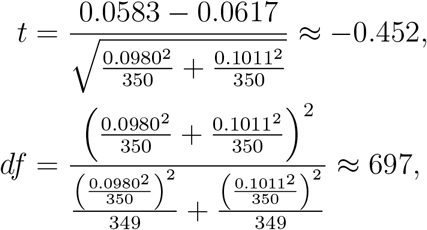

giving *p* ≈ 0.652 *> α*. Thus, across the VIPUR dataset, there is no statistically significant difference between our model’s and TM-Vec’s TM-score prediction.

### 3.3 Overall Findings

It has been demonstrated that our model’s capability for TM-score prediction is statistically indistinguishable from TM-Vec’s on both the TM-Vec dataset and the VIPUR dataset across all TM-score predictions. However, Figure 5 indicates our model is slightly more consistent at predicting low or high TM-scores (TM ∈ [0, 0.4) ∪ [0.8, 1]), and slightly worse at predicting medium TM-scores (TM ∈ [0.4, 0.8]). The improved results for true TM-scores within [0.8, 1] is possibly due to the fact that the loss function utilizes combined lDDT- and TM-score loss for high TM-scores, resulting in the model better capturing structural features at higher levels of global similarity. However, it is also likely that the differences in performance at each TM-score interval are statistically insignificant. For lDDT score prediction, our model is accurate on both datasets.

## 4 Discussion

### 4.1 Limitations

While our model has promising test results, it is important to acknowledge the current methodology’s shortcomings, including limited dataset size for both training and testing.

### 4.2 Future Work

Increasing the dataset sizes, extending training to different databases (e.g. CATH), and expanding the testing to other types of mutations would validate our model’s efficacy against a wide range of cases.

Another possible line of future study is incorporating topological priors into the model. For example, one can design a new hierarchical feature extractor that directly captures protein motifs at different neighborhood sizes (e.g. using 1D or 2D convolutions [23]), based on the prior that proteins have multiple levels of structure (residue → secondary structure → domain → global fold [24]). In this way, predicting local similarity via embeddings is no longer dependent on direct sequence alignments (like lDDT scores require).

## 5 Conclusion

State-of-the-art structural alignment algorithms, most of which rely on direct superposition, are algorithmically complex and have high time complexities. Currently, no PLMs that attempt to calculate structural similarity encapsulate both global and local structural similarity, although both are important in determining protein function, homology, and more.

This study developed a framework for a sequence-based PLM, consisting of a transformer-based Siamese neural network, which produced locally- and globally-structure aware embeddings. Our model was trained to predict both TM-score, a global similarity metric, and lDDT scores, a per-residue similarity metric.

Our testing results confirmed the plausibility of our framework, as our model performed similarly in TM-score prediction when compared to highly accurate models like TM-Vec while also producing accurate lDDT scores. This dual capability makes the model a potential tool in downstream research tasks, particularly mutation analysis, where it can aid in the identification of deleterious mutations as well as the recognition of affected subdomains.

## 6 Code Availability

My data preprocessing, training, and testing scripts are all located at https://github.com/Brainana/GLASS-PRIMES.

## Acknowledgements

I would like to thank Dr. Shaojun Pei, Dr. Gil Alterovitz, and Dr. Ning Xie for their invaluable mentorship in shaping the direction of the research. I would specifically like to thank Dr. Shaojun Pei for guiding me towards a case study on local structural changes induced by mutations, and providing me with computing resources and software that aided the development of my model. I would also like to thank the MIT PRIMES program for providing the opportunity to perform this research.

## Notes

### Competing Interest Statement

The authors have declared no competing interest.

